# Framework for Denoising Monte Carlo Photon Transport Simulations Using Deep Learning

**DOI:** 10.1101/2022.01.19.477008

**Authors:** Matin Raayai Ardakani, Leiming Yu, David R. Kaeli, Qianqian Fang

**Affiliations:** Northeastern University, Department of Electrical and Computer Engineering, 360 Huntington Ave., Boston, MA, USA, 02118; Northeastern University, Department of Bioengineering, 360 Huntington Ave., Boston, MA, USA, 02118; Analogic Corporation, 8 Centennial Drive, Peabody, MA, USA, 01960

**Keywords:** Monte Carlo Method, Image Denoising, Photon Transport, Convolutional Neural Networks, Deep Learning

## Abstract

**Significance:** The Monte Carlo (MC) method is widely used as the gold-standard for modeling light propagation inside turbid media like human tissues, but combating its inherent stochastic noise requires one to simulate large number photons, resulting in high computational burdens.

**Aim:** We aim to develop an effective image denoising technique using deep learning (DL) to dramatically improve low-photon MC simulation result quality, equivalently bringing further acceleration to the MC method.

**Approach:** We have developed a cascade-network combining DnCNN with UNet, in the meantime, extended a range of established image denoising neural-network architectures, including DnCNN, UNet, DRUNet, and ResMCNet, in handling three-dimensional (3-D) MC data and compared their performances against model-based denoising algorithms. We have also developed a simple yet effective approach to create synthetic datasets that can be used to train DL based MC denoisers.

**Results:** Overall, DL based image denoising algorithms exhibit significantly higher image quality improvements over traditional model-based denoising algorithms. Among the tested DL denoisiers, our Cascade network yields a 14 - 19 dB improvement in signal-noise ratio (SNR), which is equivalent to simulating 25 × to 78 × more photons. Other DL-based methods yielded similar results, with our method performing noticeably better with low-photon inputs, and ResMCNet along with DRUNet performing better with high-photon inputs. Our Cascade network achieved the highest quality when denoising complex domains, including brain and mouse atlases.

**Conclusion:** Incorporating state-of-the-art DL denoising techniques can equivalently reduce the computation time of MC simulations by one to two orders of magnitude. Our open-source MC denoising codes and data can be freely accessed at http://mcx.space/.

## 1 Introduction

Non-ionizing photons in the near-infrared (NIR) wavelength range have many benefits in biomedical applications compared to ionizing ones such as x-ray. Because of the low energy, NIR light is relatively safe to use and can be applied more frequently; the relatively low cost and high portability of NIR devices makes them excellent candidates for addressing needs in functional assessment on the bedside or natural environments.^1^ However, the main challenge of using low energy NIR photons is the high degree of complex interactions with human tissues due to the presence of high scattering, which is much greater than that of x-rays. As a result, the success of many emerging NIR-based imaging or intervention techniques, such as diffuse optical tomography (DOT),^2^ functional near-infrared spectroscopy (fNIRS),^3^ photobiomodulation (PBM)^4^ etc, requires a quantitative understanding of such complex photon-tissue interactions via computation-based models.

The Monte Carlo method is widely regarded as the gold-standard for modeling photon propagation in turbid media,^5^ including human tissue, due to its accuracy and flexibility.^6^ It stochastically solves the general light propagation model – the radiative transfer equation (RTE) – without needing to build large simultaneously linear equations.^7^ While an approximation of RTE, the diffusion equation (DE), can be computed more efficiently using finite element (FE)-based numerical solvers,^8^ DE is known to yield problematic solutions in regions that contain low-scattering media.^9^ Besides accuracy and generality, simplicity in implementation of MC algorithms compared to other methods has not only made MC a top choice for teaching tissue-optics, and also for developing open-source modeling tools.

MC methods have attracted even greater attention in recent years as simulation speed has increased dramatically due to the broad adoptions of massively-parallel computing and graphics processing unit (GPU) architectures. The task parallel nature of MC algorithms allows it to be efficiently map to the GPU hardware.^10^ Current massively parallel MC photon propagation algorithms are capable of handling arbitrary 3-D heterogeneous domains and have achieved hundreds fold speedups compared to traditional serial simulations.^11–15^ This breakthrough in the MC algorithm allows biophotonics researchers to increasingly use it in routine data analyses, image reconstructions and hardware parameter optimizations, in addition to its traditional role of providing reference solutions in many biophotonics domains.

A remaining challenge in MC algorithm development is the presence of stochastic noise, which is inherent in the method itself. Because an MC solution is produced by computing the mean behaviors from a large number of photon packets, each consisting of a series of random samplings of the photon scattering/absorption behaviors, creating high-quality MC solutions typically requires simulations of tens to hundreds of millions of photons. This number depends heavily on the domain size, discretization resolution and tissue optical properties. This translates to longer simulation times, because the MC runtime is typically linearly related to the number of simulated photons. From our recent work,^16^ a 10-fold increase of photon number typically results in a 10 decibel (dB) (SNR) improvement in MC solutions, suggesting that MC stochastic noise is largely shot-noise bound. From this prior work, we have also observed that the MC stochastic noise is spatially varying and, in highly scattering/absorbing tissues, exhibits high dynamic range throughout the simulation domain.

To obtain high quality simulation results without increasing the number of simulated photons, signal processing techniques have been investigated to remove the stochastic noise introduced by the MC process. This procedure is commonly referred to as denoising.^16, 17^ In the past, model-based noise-adaptive filters have been proposed to address the spatially varying noise in the radiation dosage estimation context and computer graphics rendering.^18–20^ However, improvements provided by applying these filtering-based techniques have been small to moderate, creating an equivalent speedup of only 3- to 4-fold.^16^ Recent work on denoising ray-traced computer graphics, and spatially-variant noisy images in the field of computer vision, focus mainly on machine learning (ML)-based denoising methods, more specifically convolutional neural networks (CNNs).^17^ Despite their promising performance compared to traditional filters, no attempt has been made, to the best of our knowledge, to adapt denoisers designed for two-dimensional (2-D) low bit-depth image domain to high dynamic range MC fluence maps.^16, 21^ Our motivation is therefore to develop effective CNN-based denoising techniques and compare it among state-of-the-art denoisers in the context of MC photon simulations and identify their strengths compared to traditional model-based filtering techniques.

In recent years, the emergence of convolutional neural network (CNN)s has revolutionized many image-processing-centered applications, including pattern recognition, image segmentation and super-resolution. CNNs have also been explored in image denoising applications, many targeted at removing additive white Gaussian noise from natural images,^22^ and more recently, real camera noise.^23,24^ Compared to classical approaches, CNNs have also demonstrated impressive adaptiveness to handle spatially varying noise.^25,26^ In a supervised setting, given a dataset representative of media encountered in real-life simulations, CNNs have shown to better preserve sharp edges of objects without introducing significant bias compared to model-based methods.^22, 27, 28^ Finally, due to extensive efforts over the past decade to accelerate CNNs on GPUs, modern implementations of CNN libraries can readily take advantages of GPU hardware to achieve high computational speed compared to traditional methods. Nonetheless, there has not been a systematic study to quantify CNN image denoiser performance over MC photon transport simulation images, either in 2-D or 3-D domains.

The contributions of this work are the following. First, we have developed a simple generative model that uses the Monte Carlo eXtreme (MCX)^12^ software to create a synthetic dataset suited for supervised training of an image denoiser, providing ample opportunities for learning its underlying noise structure. Secondly, we have developed and characterized a novel spatial-domain CNN model that cascades DnCNN^26^ (an effective global denoiser) and UNet^29^ (an effective local denoiser). Thirdly, we have adapted and quantitatively compared a range of state-of-the-art image denoising networks, including DnCNN,^26^ UNet,^29^ DRUNet,^28^ deep residual-learning for denoising MC renderings^30^ (referred to as ResMCNet hereinafter), as well as our cascaded denoiser, in the context of denoising 3-D MC simulations. We assess these methods using a number of evaluation metrics, including mean-squared error (MSE) and structural similarity index measure (SSIM). For simplicity, other DL-based denoising methods that do not operate in the spatial domain,^31,32^ or require specialized knowledge from their target domain,^33^ are not investigated here and left for future work. Lastly, a range of challenges encountered during the development of our approach are also discussed, providing guidance to future work in this area.

## 2 Methods

### 2.1 Training Dataset Overview

To train and evaluate CNN denoisers in a supervised fashion, a series of datasets were generated that provided one-to-one mappings between “noisy” and “clean” simulations. The training dataset was created using our MCX software package,^12^ in which simulations of a range of configurations with different photon levels were included. The 3-D fluence maps generated from the highest number of photons were treated as “clean” data, and the rest were regarded as noisy. For this work, all configurations were simulated with photon numbers between 10^5^ and 10^9^ with an increment of 10-fold. Simulations with 10^9^ photons were selected as the “ground-truth”, since they provide the closest estimate to the noise-free solutions. Therefore, the CNN denoisers are tasked to learn a mapping between simulations with photon numbers lower than 10^9^ to results simulated with 10^9^ photons.

#### 2.1.1 Generation of Training and Validation Datasets

To efficiently generate a large and comprehensive corpus of representative MC training data, first a volume generation scheme was designed. In such scheme, arbitrary-shaped and sized polyhedrons and random 3-D American standard code for information interchange (ASCII) characters with arbitrary sizes are randomly placed inside a homogeneous background domain with random optical properties. Using combinations of ASCII characters and polyhedrons produces a wide variety of complex shapes, while keeping the data generation process efficient. A diagram showing the detailed steps for creating a random simulation domain for generating training data is shown in Fig. 1.

**Fig 1:**
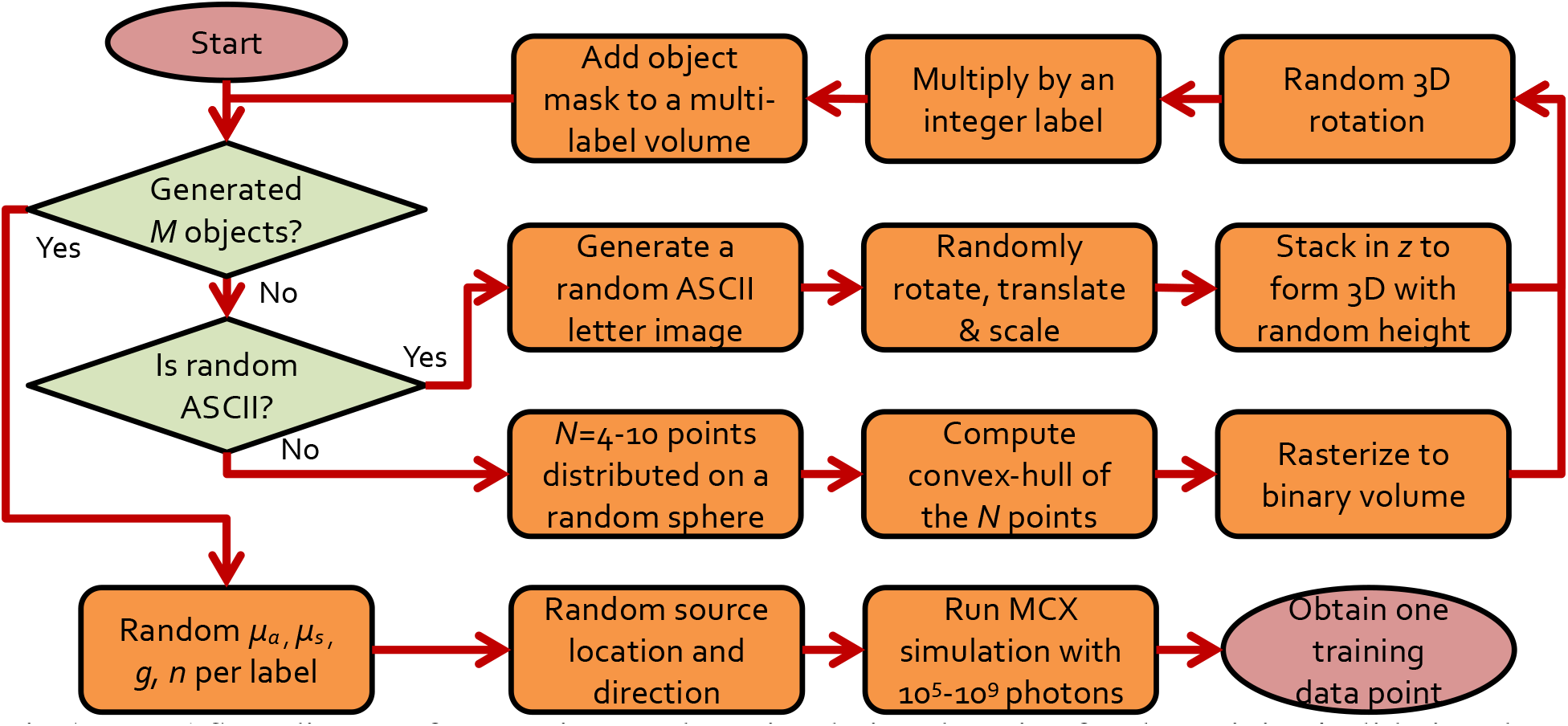
Workflow diagram for creating random simulation domains for the training/validation data.

Specifically, a random number (M = 0 to 4) of randomly generated shapes, either in the form of 3-D polyhedrons, or 3-D ASCII letters, are first created as binary masks, with the same size as the target volume. Then, the binary mask is multiplied by a label – a unique identification number assigned to each object – and subsequently accumulated in a final volume, where voxels marked with the same label belong to the same shape. In the process of accumulation and generation of binary masks for each shape, if two or more objects intersect, this process creates new inclusions for the overlapping regions. We generated all training datasets on a 64 × 64 × 64 (in 1 mm^3^ isotropic voxels) domains, while the datasets for validation were 128 × 128 × 128 voxels. This allows us to observe the scalability of the networks to volume sizes different than the training dataset. A total of 1,500 random domains were generated for training and 500 random domains for the “validation”. During training, the average global metrics (explained in Section 2.4.1) of the model computed over the validation dataset were saved over single epoch intervals. At the end of the training, the model with the best overall metrics was selected as the final result.

To create random 3-D polyhedrons, a number of points (*N* = 4 to 10) are determined on a sphere of random location and radius using the algorithm provided by Deserno *et al*.^34^ The convex-hull of the point set is computed and randomly rotated and translated in 3-D. This convex-hull is subsequently rasterized into a binary mask.

For ASCII character inclusions, first, a random character in either lower or upper cases of English alphabet is selected. A random font size is chosen from a specified range, and the letter is rendered/rasterized in a 2-D image with a random rotation angle and position. This binary 2-D mask is further stacked with a random thickness to form a 3-D inclusion. Finally, a 3-D random rotation/translation is applied to the 3-D ASCII character inclusion.

After generating a random volume, a random simulation configuration is generated to enable simulations with MCX. This includes determining the optical properties, including absorption (*μ_a_*), scattering (*μ_s_*) coefficients, anisotropy (*g*) and refractive index (*n*), for each of the label inside the generated volume, as well as the light source position and launch direction for the simulation. For the training and validation datasets, only isotropic sources were used for simplicity. The source is randomly positioned inside the domain.

The random optical properties are determined in ranges relevant to those of biological tissues, including: 1) *μ_a_* = |*N*(0.01; 0.05) | mm^-1^ where *N*(*μ*; **σ**) is a normal distribution with mean *μ* and standard deviation **σ**), 2) *g* is uniform random variable between 0.9 and 1.0, 3) 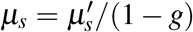 where the reduced scattering coefficient 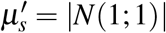 mm^−1^, and 4) *n* is a uniformly distributed random variable between 1 and 10. For all data, we simulate the continuous-wave (CW) fluence for a time-gate length randomly selected between 0.1 and 1 ns with a 0.1 ns step size. Each simulation uses a random seed. In Fig. 2, we show a number of image slices (log-10 scale) from 3-D simulation samples ranging from homogeneous domains to heterogeneous domains containing multiple polyhedral or letter-shaped inclusions.

**Fig 2:**
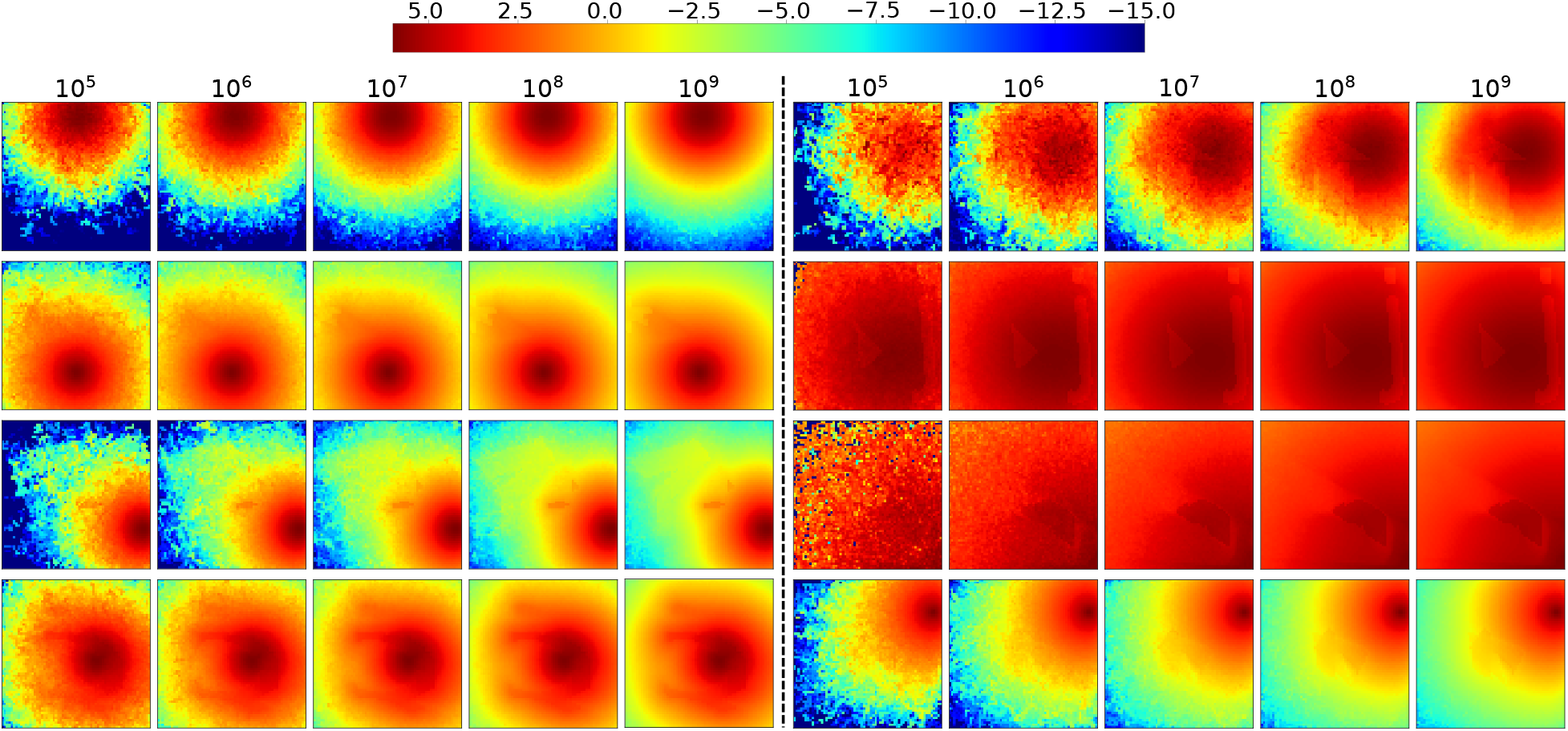
Sample MC fluence images (slices from 3-D volumes) generated for CNN training.

#### 2.1.2 Data Augmentation

To increase the diversity of the generated dataset and avoid overfitting, data augmentation^35^ was used. Our data augmentation consisted of 90-degree rotation and flipping. Each transformation was applied independently over a randomly selected axis. Transforms were identically applied to both inputs and labels of the training data. Both transforms were randomly selected and applied, with a probability of 0.7. This on-the-fly strategy multiplied the data encountered by the models during training by 256, without performing any time-consuming MC simulation.

#### 2.1.3 Test Datasets

Three previously used standard benchmarks^16^ – (B1) a 100 × 100 × 100 mm^3^ homogeneous cube with a 1 mm voxel size, (B2) same cubic domain with a 40 × 40 × 40 mm^3^ cubic absorber and (B3) same cubic domain with a refractive inclusion were employed to characterize and compare the performance of various denoising methods. The optical properties for the background medium, the absorbing and refractive inclusions can be found in Section 3 of our previous work.^16^ Each of the benchmarks were simulated with 100 repetitions using different random seeds. Additionally, the Colin27^12, 36^ atlas (B4), Digimouse^37^ atlas (B5) and USC 19.5^38^ atlas (B6) from the Neurodevelopmental MRI database^39^ were selected as examples of complex simulation domains to test our trained MC denoisers.

### 2.2 Pre-processing of Monte Carlo Data

Many of the reported DL denoising techniques were developed to process natural images of limited bit-depth that usually do not present the high dynamic range as in MC fluence maps. To allow the CNNs to better recognize and process unique MC image features and avoid difficulties due to limited precision, we applied the below transformation to the fluence images before training or inference

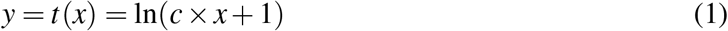

where *x* is the MC fluence map, *c* is a user-defined constant, and the output *y* serves as the input to the CNN. This transformation serves two purposes. First, it compresses the floating-point fluence values to a limited range while equalizing image features across the domain. Secondly, it compensates for the exponential decay of light in lossy media and reveals image contracts that are relevant to the shapes/locations of the inclusions, assisting the CNN to learn the features and mappings. The addition of 1 in Eq. 1 ensures that *t*(*x*) does not contain negative values. An inverse transform *t*^−1^ (y′) = (*e^y′^* – 1)/c is applied to the output of the CNN (*y′*) to undo the effect of this transform.

Moreover, when training a CNN on 8-bit natural image data, a common practice is to divide the pixel values by the maximum value possible (i.e., 255) to normalize the data. From our tests, applying such operation on floating-point fluence maps resulted in unstable training, therefore our training data were neither quantized nor normalized.

Additionally, due to limited data precision, we noticed that all tested CNN denoisers exhibit reduced denoising image quality when processing voxel values (before log-transformation) that are smaller than an empirical threshold of 0.03. To address this issue and permit a wider input dynamic range, two separate copies of the fluence maps were denoised during inference – the first copy was denoised with *c* set to 1, and the second one with *c* set to 10^7^. The final image is obtained by merging both denoised outputs: voxels that originally had fluence value larger than 0.03 retrieve the denoised values from the first output and the rest are obtained from the second output. This variable-gain approach allowed us to process MC fluence images containing both high and low floating point values.

### 2.3 A Cascaded MC Denoising Network that Combines DnCNN and UNet Networks

In this work, we designed a cascaded CNN denoiser specifically optimized for denoising our 3-D MC fluence maps by combining two existing CNN denoisers: a DnCNN denoiser is known to be effective for removing global or spatially-invariant noise, especially additive white gaussian noise (AWGN), without any prior information,^26^ while a UNet denoiser is known to remove local noise that is spatially-variant.^28^ Therefore, in our cascaded DnCNN/UNet architecture, referred to as “Cascade” hereinafter, the CNN first learns the global noise of an MC fluence image and attempts to remove it. The remaining spatially-variant noise can then be captured and removed using a UNet. In both stages, the noise is learned in the residual space, meaning that instead of mapping a noisy input to a clean output directly, the network maps the noisy input to a noise map and then subtracts it from the input to extract the clean image.

### 2.4 Denoising Performance Metrics

#### 2.4.1 Global Performance Metrics

The global resemblance between the denoised volume and the ground-truth (in this case, simulations with 10^9^ photons) can be used to measure the performance of a denoiser. A number of metrics measuring such similarity have been used by others to evaluate image restoration networks or measure convergence.^21, 26, 40, 41^ Typically, these metrics are defined for 2-D images; in this work, we have extended the definitions to apply to 3-D fluence maps.

The most commonly used objective functions for denoising networks are the mean least squared error (*L*_2_) and mean absolute error (*L*_1_):

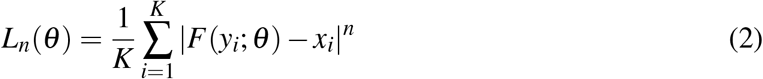

where *K* is the number of noisy-clean fluence map pairs sampled from the dataset, referred to as the “mini-batch” size, *θ* contains all parameters of the network, *F* denotes the network itself, *n* is either 1 or 2, and (*x_i_, y_i_*) denotes the i-th noisy-clean pair of data in the mini-batch. These error metrics are widely used in supervised denoising networks, including DnCNN, DRUNet and ResMCNet models, as well as several other studies.^25, 26, 28, 30, 42^ *L*_1_ and *L*_2_ may have different convergence properties.^40^ The *L*_1_ loss has gained more popularity in the DL community, due to it’s good performance and low computational costs.^30,40^ For this work, however, to penalize large errors more, the *L*_2_ loss was used instead to train the networks.

In contrast to *L_n_* distances, SSIM^43^ provides a perceptually-motivated measure that emulates human visual perception for images. The SSIM for a pixel in an image is be defined as:

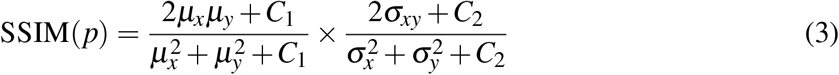

where *μ_x_* and *σ_x_* are the mean and standard deviation of the image *x*, respectively, and *σ_xy_* is the co-variance of images *x* and *y*. The statistics are calculated locally by convolving both volumes with a 2-D Gaussian filter with *σ_G_* = 5. Small constants *C*_1_ and *C*_2_ are used to avoid division by zero. The SSIM value of two images is the average SSIM computed across all pixels, with a value of 1 suggesting the two images are identical, and a value of 0 suggesting the two images are not correlated. This definition can also be applied to 3-D fluence maps by using a 3-D Gaussian kernel to calculate neighborhood statistics.

Another metric, peak signal-to-noise ratio (PSNR), measures the ratio between the maximum power of a signal and the power of the noise.^44^ The PSNR for two volumes *x* and *y* is expressed as:

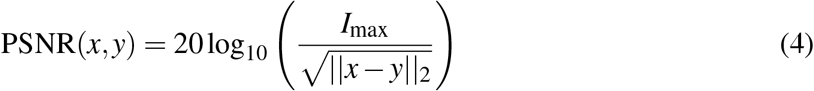

Larger PSNR values indicate smaller *L*_2_ distances between volumes. The *I_max_* value is the maximum value a voxel can have in a fluence map after the transformation in Eq. 1. Therefore, in this work, we set *I_max_* to 40.

#### 2.4.2 Local Performance Metrics

A number of locally (voxel-bound) defined performance metrics have been used in our previous MC denoising work.^16^ The SNR of the denoised volumes for each voxel measures the efficacy of the denoiser of spatially adaptive noise. For a simulation running *k* photons, we first run multiple (*N* = 100) independently-seeded MC simulations and compute SNR in dB with

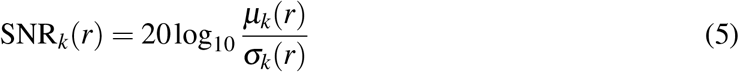

where *μ_k_* and *σ_k_* are the mean and standard deviation of voxel values at location *r* across all repetitions. The average SNR difference before and after applying the denoising filter, ΔSNR, is subsequently calculated along selected regions-of-interest.

Our previous work^16^ suggests that the noise in MC images largely follows the shot-noise model; therefore, increasing the simulated photon number by a factor of 10 results in ~ 10 dB improvement in SNR on average. We have previously proposed a photon number multiplier16 *M_F_* to measure equivalent acceleration using the average SNR improvement ΔSNR

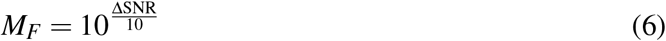

A ΔSNR = 20 dB gives *M_F_* = 100, suggesting that the denoised result is equivalent to a simulation with 100 times of the originally simulated photon number, thus is equivalent to accelerating the simulation by a factor of 100 if the denoising run-time is ignored.

### 2.5 Implementation Details

#### 2.5.1 BM4D and ANLM

Block-matching 4-D collaborative filtering (BM4D) and our GPU-accelerated adaptive non-local means (ANLM)^16^ are used as representative state-of-the-art model-based denoisers and used to compare against CNN based denoisers. For BM4D, a Python interface developed based on the filter described by Makinen *et al*.^45^ was used, whereas for the ANLM filter, a MATLAB function developed previously by our group^16^ was used.

#### 2.5.2 CNN Training Details

All CNN denoising networks were re-implemented for handling 3-D data using an open-source DL framework, PyTorch.^46^ For most of the studied CNN denoisers, our implementations largely follow their originally published specifications, while replacing the 2-D layers with their 3-D variants. Small adjustments were made. For UNet, for example, 3-D batch normalization layers were introduced in-between the 3-D convolution, the convolution transpose, and the pooling layers to address the covariance shift problem.^47^ Additionally, we have simplified ResMCNet by removing the auxiliary features needed for computer graphics renderings purposes, making the kernel size of the first layer 3 instead of 7.

All networks in this study were trained for 1,500 epochs on a single NVIDIA DGX node equipped with 8 NVIDIA A100 GPUs, each with 40 GB of memory and NVLink 2.0 connection. Leveraging the PyTorch scaling wrapper, PyTorch Lightning^48^ was used to simplify the implementation process. We need high performance hardware since a forward propagation of the CNN for a 64 × 64 × 64 voxelated volume requires around 6 GB of GPU memory; to use a batch size of 4 per GPU (i.e. processing 4 data pairs in parallel), at least 24 GB of memory is necessary. Furthermore, using all 8 GPUs in parallel combined with the high-speed NVLink connection reduces the average training time from 10 days (on a single A100 GPU) to 24 hours for each network tested – the Cascade and DRUNet usually require longer training time compared to those of DnCNN and UNet.

The networks were all trained using the “Adam with weight decay regularization (AdamW)” optimizer,^49^ with a weight decay of 0.0001 for the parameters in all layers, except for the batch normalization parameters and bias parameters. The learning rate was scheduled with a “cosine annealing” learning rate,^50^ using 1,000 linear warm-up mini-batch iterations to added learning stability.^51^ A batch-size of 4 per GPU was selected to maximize the effective use of GPU memory resources. The base learning rate was set to 0.0001. The gradient clipping value was set to 2 for batch normalization layers, and 1 for other layers to avoid exploding gradients and faster training.^52^ The optimization, data augmentation, and configuration sections the codebase for this work were inspired by the open-source PyTorch Connectomics package^53^ for easier prototyping of the trained models.

## 3 Results

### 3.1 Denoising Performance

In Fig. 4, we visually compare the fluence maps before and after denoising for each tested denosier and photon number (10^5^ to 10^8^) for 3 standard benchmarks^16^ (B1, B2 and B3). Table 1 summarizes the global metrics derived from the outputs of each denoiser; computed local metrics including mean ΔSNR and *M_F_* are reported in Table 3. Each entry in both tables is averaged from 100 independently-seeded repeated simulations. In both tables, the best-performing metrics are highlighted in bold. Similarly, a visual comparison between those from more complex domains, including Colin27, Digimouse, and USC-19.5 atlases, are shown in Fig. 6 and the corresponding global metrics are summarized in Table 2. Due to limited space, in Fig. 6, we only show representative images with 10^5^ and 10^7^ photons, and removed DnCNN and BM4D due to their relatively poor performances.

**Fig 3:**
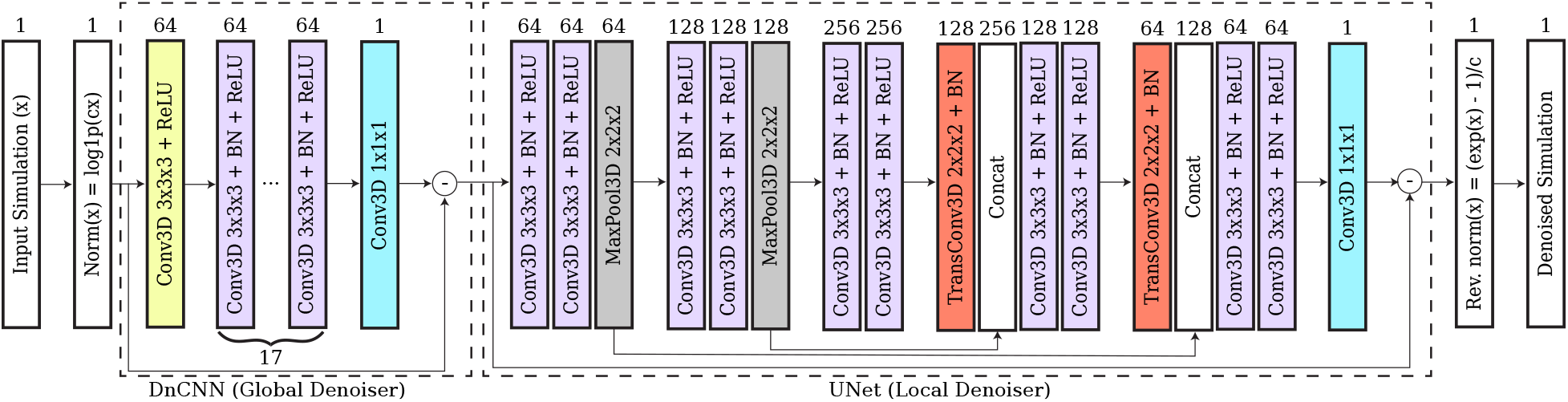
Overview of the cascaded DnCNN + UNet architecture. Each block in the dashed squares represents a group of CNN layers that are applied sequentially. The number on the square block indicates the number of channels for the respective output tensor. Conv3D, TransConv3D, and BN stand for 3D convolution, 3D transposed convolution, and batch normalization layers, respectively. PyTorch function `log1p(cx)` is a stable implementation of function ln(*cx* + 1).

**Fig 4:**
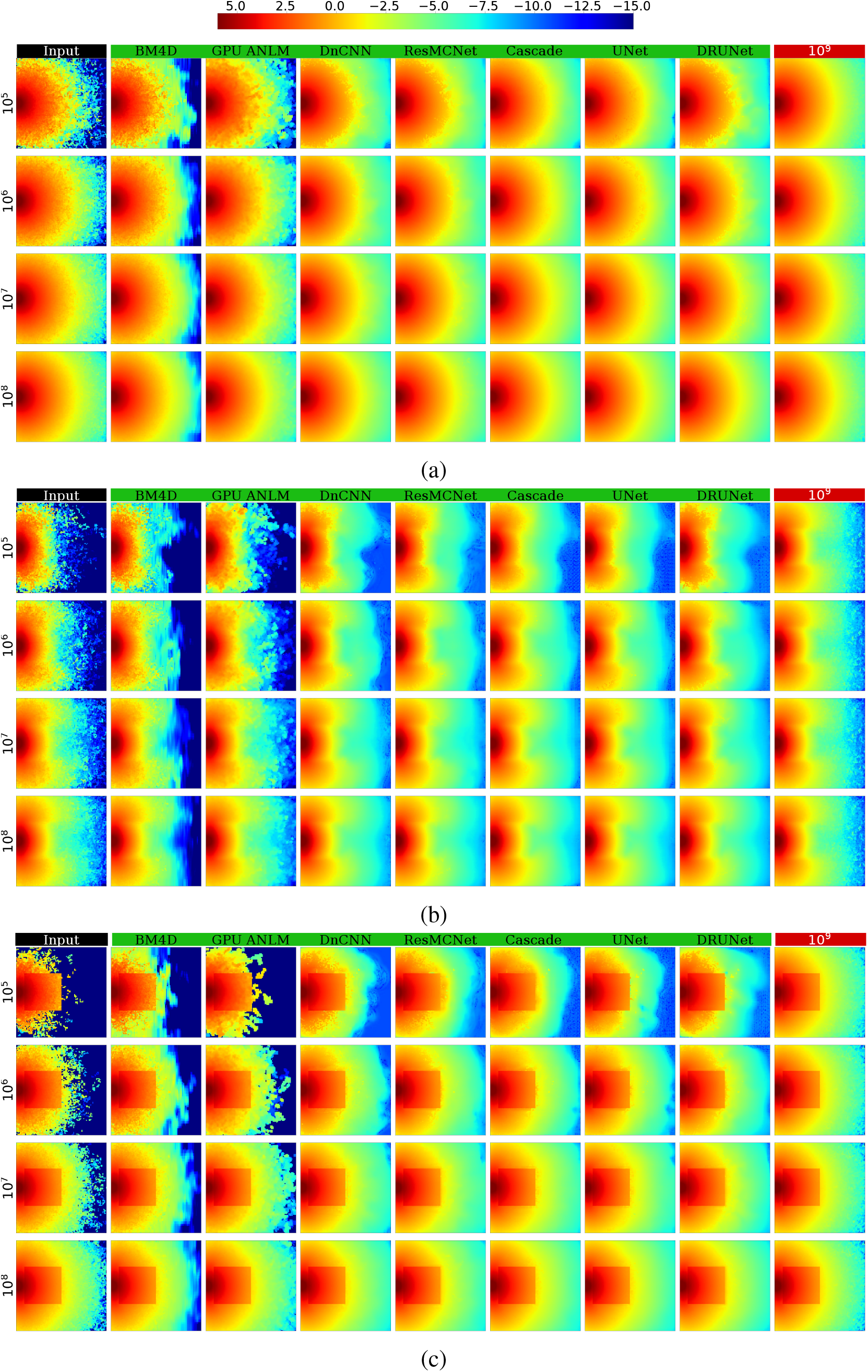
Comparisons between various denoisers in 3 benchmarks: (a) a homogeneous cube, and the same cube containing inclusions with (b) absorption and (c) refractive-index contrasts.

**Table 1:** Average global metrics derived from 3 basic benchmarks: (B1) a homogeneous cube, and the same cube with (B2) an absorption and (B3) refractive index inclusion; each data point was averaged over 100 repetitions. The best performing models are highlighted in bold.

| Metric |  | Cascade |  |  | UNet |  |  | ResMCNet |  |  | DnCNN |  |  |
| --- | --- | --- | --- | --- | --- | --- | --- | --- | --- | --- | --- | --- | --- |
|  |  | B1 | B2 | B3 | B1 | B2 | B3 | B1 | B2 | B3 | B1 | B2 | B3 |
| MSE | 10<sup>5</sup> | **0.0031** | **0.0071** | **0.0192** | 0.0050 | 0.0087 | 0.0210 | 0.0066 | 0.0114 | 0.0224 | 0.0121 | 0.0245 | 0.0507 |
|  | 10<sup>6</sup> | 0.0005 | **0.0009** | 0.0036 | 0.0012 | 0.0017 | 0.0047 | **0.0004** | 0.0010 | **0.0028** | 0.0008 | 0.0017 | 0.0044 |
|  | 10<sup>7</sup> | 0.0001 | 0.0002 | **0.0007** | 0.0001 | 0.0003 | 0.0012 | 0.0001 | 0.0002 | 0.0009 | 0.0002 | 0.0003 | 0.0008 |
| SSIM | 10<sup>5</sup> | **0.9472** | **0.9063** | **0.8663** | 0.9194 | 0.8776 | 0.8343 | 0.8741 | 0.8731 | 0.8345 | 0.8154 | 0.8131 | 0.7679 |
|  | 10<sup>6</sup> | **0.9690** | **0.9769** | **0.9670** | 0.9117 | 0.9520 | 0.9541 | 0.9680 | 0.9600 | 0.9472 | 0.9610 | 0.9392 | 0.9174 |
|  | 10<sup>7</sup> | 0.9891 | 0.9839 | 0.9817 | 0.9757 | 0.9580 | 0.9594 | 0.9930 | **0.9893** | **0.9851** | 0.9924 | 0.9876 | 0.9812 |
| PSNR | 10<sup>5</sup> | **57.1445** | **53.5681** | **49.2250** | 55.0538 | 52.7036 | 48.8291 | 53.8926 | 51.5285 | 48.5567 | 51.2112 | 48.1769 | 44.9994 |
|  | 10<sup>6</sup> | 65.4709 | **62.2454** | 56.5081 | 61.3193 | 59.6665 | 55.2916 | **65.6337** | 61.9344 | **57.5427** | 62.9706 | 59.7418 | 55.6253 |
|  | 10<sup>7</sup> | 70.9347 | 68.5446 | **63.4404** | 70.5365 | 67.4766 | 61.1240 | 71.3430 | 69.4779 | 62.3238 | 69.5030 | 67.9231 | 62.9773 |
|  |  | DRUNet |  |  | GPU-ANLM |  |  | BM4D |  |  |  |  |  |
|  |  | B1 | B2 | B3 | B1 | B2 | B3 | B1 | B2 | B3 |  |  |  |
| MSE | 10<sup>5</sup> | 0.0105 | 0.0244 | 0.0400 | 0.0830 | 0.0752 | 0.1009 | 0.2223 | 0.1826 | 0.4005 |  |  |  |
|  | 10<sup>6</sup> | 0.0005 | 0.0013 | 0.0037 | 0.0130 | 0.0132 | 0.0163 | 0.0395 | 0.0383 | 0.3475 |  |  |  |
|  | 10<sup>7</sup> | **0.0001** | **0.0002** | 0.0011 | 0.0017 | 0.0018 | 0.0022 | 0.0051 | 0.0056 | 0.3366 |  |  |  |
| SSIM | 10<sup>5</sup> | 0.8223 | 0.8115 | 0.7728 | 0.6745 | 0.7579 | 0.7092 | 0.5676 | 0.7102 | 0.5802 |  |  |  |
|  | 10<sup>6</sup> | 0.9578 | 0.9355 | 0.9227 | 0.8463 | 0.8555 | 0.8304 | 0.7174 | 0.7790 | 0.6130 |  |  |  |
|  | 10<sup>7</sup> | **0.9933** | 0.9877 | 0.9829 | 0.9627 | 0.9521 | 0.9459 | 0.8945 | 0.8862 | 0.6605 |  |  |  |
| PSNR | 10<sup>5</sup> | 51.8499 | 48.1884 | 46.0353 | 42.8553 | 43.2828 | 42.0060 | 38.5718 | 39.4263 | 36.0164 |  |  |  |
|  | 10<sup>6</sup> | 65.1219 | 61.0106 | 56.3168 | 50.8978 | 50.8340 | 49.9111 | 46.0782 | 46.2030 | 36.6318 |  |  |  |
|  | 10<sup>7</sup> | **71.6660** | **69.6645** | 61.6879 | 59.8368 | 59.4459 | 58.6816 | 54.9554 | 54.5706 | 36.7706 |  |  |  |

**Table 2:** Average global metrics derived from 3 complex benchmarks: (B4) Colin27, (B5) Digimouse and (B6) USC-195 atlases; each data point was averaged over 100 repetitions. The best performing models are highlighted in bold.

| Metric |  | Cascade |  |  | UNet |  |  | ResMCNet |  |  | DnCNN |  |  |
| --- | --- | --- | --- | --- | --- | --- | --- | --- | --- | --- | --- | --- | --- |
|  |  | B4 | B5 | B6 | B4 | B5 | B6 | B4 | B5 | B6 | B4 | B5 | B6 |
| MSE | $10^5$ | **0.0121** | **0.0701** | 0.0192 | 0.0121 | 0.0741 | **0.0179** | 0.0134 | 0.0878 | 0.0197 | 0.0162 | 0.1314 | 0.0243 |
| | $10^6$ | **0.0019** | **0.0072** | **0.0025** | 0.0020 | 0.0073 | 0.0027 | 0.0022 | 0.0097 | 0.0028 | 0.0022 | 0.0115 | 0.0028 |
| | $10^7$ | **0.0005** | **0.0011** | **0.0007** | 0.0006 | 0.0013 | 0.0007 | 0.0009 | 0.0027 | 0.0011 | 0.0006 | 0.0014 | 0.0007 |
| SSIM | $10^5$ | 0.9325 | **0.8893** | 0.9193 | 0.9311 | 0.8884 | 0.9187 | **0.9333** | 0.8795 | **0.9213** | 0.9221 | 0.8676 | 0.9111 |
| | $10^6$ | **0.9699** | **0.9548** | **0.9588** | 0.9668 | 0.9464 | 0.9574 | 0.9670 | 0.9373 | 0.9580 | 0.9608 | 0.9291 | 0.9508 |
| | $10^7$ | **0.9875** | **0.9800** | 0.9825 | 0.9833 | 0.9702 | 0.9807 | 0.9870 | 0.9765 | **0.9825** | 0.9846 | 0.9737 | 0.9791 |
| PSNR | $10^5$ | **51.2382** | **43.5914** | 49.2199 | 51.2260 | 43.3460 | **49.5335** | 50.7901 | 42.6124 | 49.1071 | 49.9522 | 40.8572 | 48.1905 |
| | $10^6$ | **59.3005** | **53.4987** | **58.0114** | 58.9310 | 53.4007 | 57.7991 | 58.5947 | 52.1780 | 57.6404 | 58.5953 | 51.4205 | 57.5059 |
| | $10^7$ | **64.8770** | **61.6786** | **63.8532** | 64.4113 | 60.8132 | 63.6941 | 62.6247 | 57.7291 | 61.8291 | 64.2162 | 60.6277 | 63.5614 |
|  |  | DRUNet |  |  | GPU-ANLM |  |  | BM4D |  |  |  |  |  |
|  |  | B4 | B5 | B6 | B4 | B5 | B6 | B4 | B5 | B6 |  |  |  |
| MSE | $10^5$ | 0.0159 | 0.1246 | 0.0233 | 0.0667 | 0.3393 | 0.0724 | 0.0807 | 0.4070 | 0.1006 | | | |
| | $10^6$ | 0.0023 | 0.0119 | 0.0028 | 0.0407 | 0.1860 | 0.0383 | 0.0187 | 0.0910 | 0.0242 | | | |
| | $10^7$ | 0.0007 | 0.0020 | 0.0007 | 0.0325 | 0.1244 | 0.0283 | 0.0032 | 0.0160 | 0.0044 | | | |
| SSIM | $10^5$ | 0.9213 | 0.8747 | 0.9118 | 0.8995 | 0.8379 | 0.8941 | 0.8891 | 0.8298 | 0.8853 | | | |
| | $10^6$ | 0.9583 | 0.9307 | 0.9492 | 0.9290 | 0.8756 | 0.9210 | 0.9134 | 0.8693 | 0.9068 | | | |
| | $10^7$ | 0.9843 | 0.9734 | 0.9792 | 0.9615 | 0.9214 | 0.9542 | 0.9467 | 0.9164 | 0.9377 | | | |
| PSNR | $10^5$ | 50.0399 | 41.0909 | 48.3797 | 43.8036 | 36.7362 | 43.4453 | 42.9745 | 35.9452 | 42.0139 | | | |
| | $10^6$ | 58.5107 | 51.2805 | 57.5798 | 45.9432 | 39.3454 | 46.2047 | 49.3282 | 42.4529 | 48.1976 | | | |
| | $10^7$ | 63.9421 | 59.1151 | 63.7079 | 46.9164 | 41.0927 | 47.5314 | 56.9748 | 50.0000 | 55.6500 | | | |

From the denoised images shown in Fig. 4, we can first confirm that all CNN based denoisers show noise-adaptive capability similar to ANLM and BM4D – they apply higher level of smoothing in noisy areas within low-photon regions, and apply little smoothing in areas with sufficient photon fluence. From Fig. 4c, we can also observe that all CNN denoisers show edge-preservation capability, again similar to ANLM and BM4D. Both noise-adaptiveness and edge-preservation are considered desirable for an MC denoiser.^16^ Because all CNN networks were trained on images of 64 × 64 × 64 voxels while all 3 benchmarks shown in Fig. 4 are 100 × 100 × 100 voxel domains, these results clearly suggest that our trained networks can be directly applied to image domain sizes different from the training domain size.

By visually inspecting and comparing the denoised images in Figs. 4 and 6, we observed that all CNN based methods appear to achieve significantly better results compared to model-based denoising methods (BM4D and GPU ANLM); such difference is even more pronounced in low-photon simulations (10^5^ and 10^6^ photons). Although the CNN denoisers were trained on shapes with less complexity, the images in Fig. 6 indicate that they are clearly capable of denoising novel structures that are significantly complex, yielding results that are close to the respective groundtruth images. However, we also observe that the denoiser’s ability to recover fluence maps varies depending on the photon level in the input data – in areas where photons are sparse, the denoisers understandably create distortions that deviate from the ground-truth. Nevertheless, these distorted recovered areas are still significantly better than the input in the same area without denoising.

To confirm that CNN denoisers can produce unbiased images, the means and SNRs from benchmarks B1, B2 and B3 along the line *x* = 50 and *y* = 50 were calculated and plotted in Fig. 5. For brevity, we only report the results from the Cascade network as representative of all CNN methods in this plot. These plots confirm that the Cascade method does not alter the mean fluence of the simulations over the plotted cross section, while providing a consistent SNR improvement across a wide range of photon numbers. It also demonstrates that the adaptiveness of CNN denoisers, that SNR improvement starts to decline in areas with high fluence value (thus lower noise due to shot-noise). The ~12 dB SNR improvement shown by denoising simulations with 10^9^ photons (purple dotted lines over purple solid lines in the SNR plots) indicate that the Cascade denoiser is capable of further enhancing image quality even it was not trained using simulations with more than 10^9^ photons. Such SNR improvement is not as high as that reported from low-photon simulations, yet, it is still significantly higher than the best SNR improvement produced using GPU ANLM denoiser (dashed lines) of all tested photon numbers.

**Fig 5:**
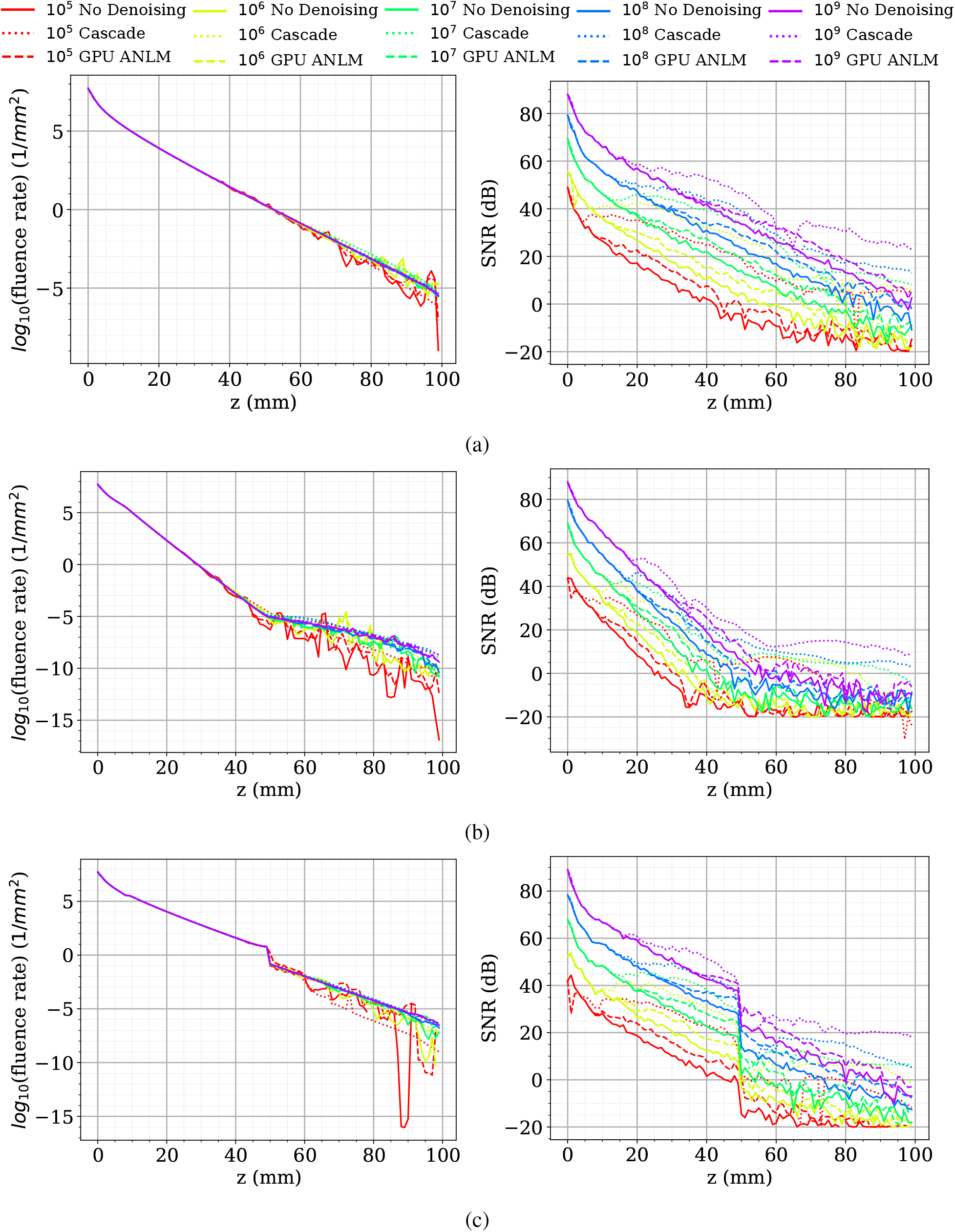
Plots of the means (left) and SNRs (right) before (solid) and after denoising using Cascade network (dotted) and GPU-ANLM (dashed) in 3 benchmarks (a) B1, (b) B2 and (c) B3 along a cross-section.

**Fig 6:**
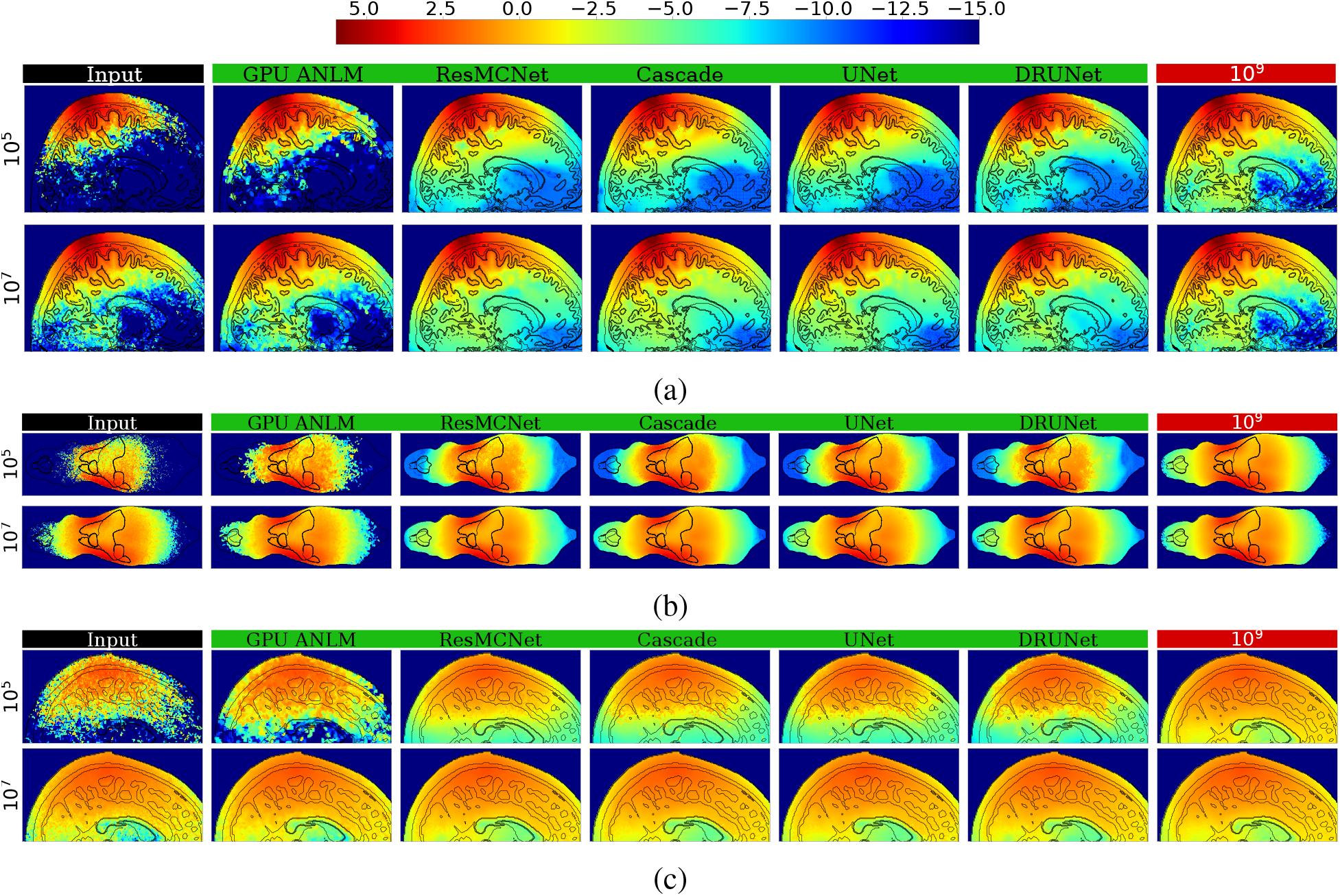
Comparisons between various denoisers in 3 complex benchmarks: (a) a Colin27, (b) Digimouse and (c) USC-195 atlases.

Our earlier observation that most CNN based denoisers outperform model-based denoisers (GPU ANLM and BM4D) is also strongly evident by both the global metrics reported in Table 1 and local metrics reported in Table 3. Among all tested CNN filters, the Cascade network offers the highest performance in all tests with 10^5^ photons, and comes close to the best performer – ResMCNet – among the 10^6^ test sets. Among the 10^7^ photon levels, DRUNet is a strong performer, with ResMCNet and Cascade coming close or surpassing it in some cases. Among the real-world complex domain benchmarks shown in Table 2, Cascade reports the best performance in almost all cases with UNet performing slightly better on USC-195 with 10^5^ photons and ResMCNet giving better SSIM results.

**Table 3:**
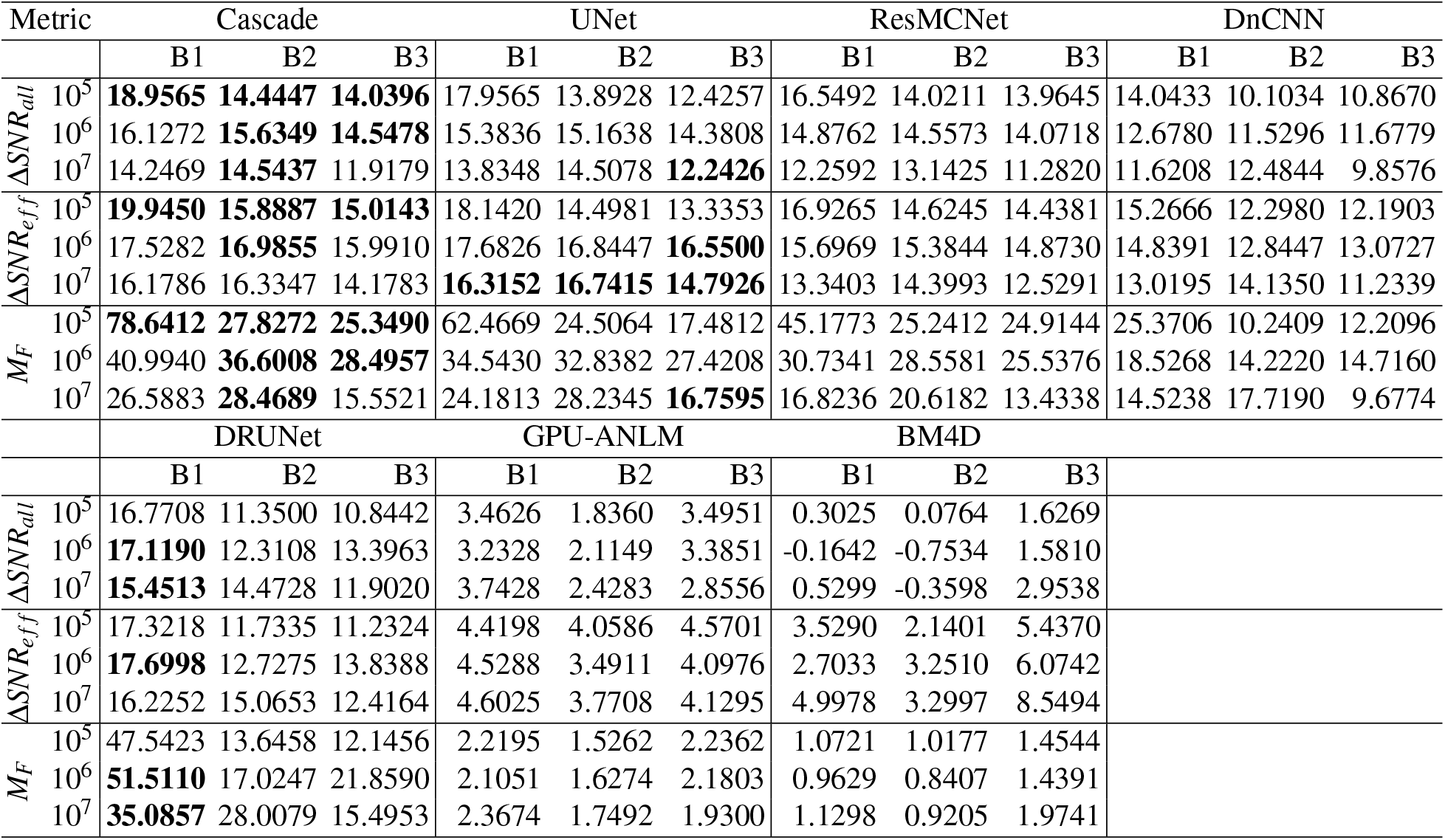
Overall average SNR improvements (ΔSNR_*all*_ in dB) and those (ΔSNR_*eff*_) in the effective region (where ΔSNR > 0.5 dB) as well as the photon number multipliers (*M_F_*) in the 3 basic benchmarks (B1-B3).

From Table 3, we can observe that all CNN based denoisers appear to offer a 5-8 fold improvement in SNR enhancement compared to our previously reported model-based GPU ANLM filter;^16^ our Cascade network reports an overall SNR improvement between 14 to 19 dB across different benchmarks and photon numbers. This is equivalent to running 25× to 35× more photons in heterogeneous domains, and nearly 80× more photons for the homogeneous benchmark (B1). In other words, applying our Cascade network for an MC solution with 10^5^ photons can obtain a result that is equivalent to running ~ 2.5 × 10^6^ photons. In fact, except for DnCNN, the majority of our tested CNN based denoisers can achieve a similar level of performance.

### 3.2 Assessing Equivalent Speed-up Enabled by Image Denoising

In Table 4, we report the average runtimes (in seconds) of MC simulation and denoising (i.e. inference for CNN denoisiers). Each test case runs on a single NVIDIA A100 with 40 GBs of memory with over 100 trials, and the time needed to transfer data between the host and the GPU is included. As we mentioned in Section 2.2, to obtain every denoised image, we apply CNN inference twice to handle high dynamic range in the input data.

**Table 4:**
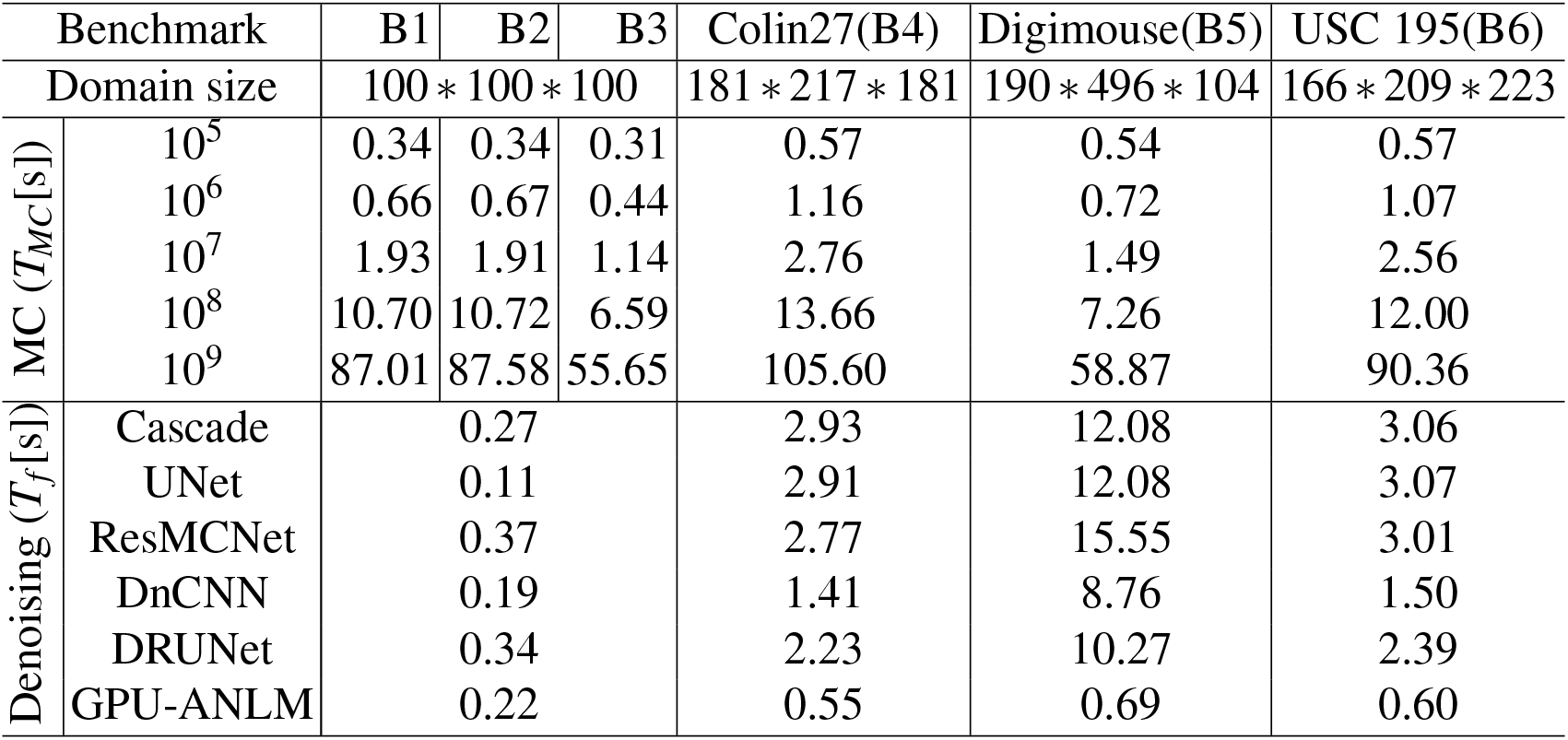
Average runtimes (in seconds) for MC forward simulations (*T_MC_*) and denoising (*T_f_*) across all benchmarks, measured on an NVIDIA A100 GPU. The runtimes include memory transfer operations.

Table 3 suggests that on average, about a 20 to 30 photon multiplier (*M_F_*) is to be expected for most CNN denoisers, meaning the denoised simulations will have 20 to 30 times more photons than its input. Therefore, our goal is to identify cases where the sum of the runtime of the baseline MC simulation running on *N* photons, *T_MC_(N)*, and that of the denoiser (*T_f_*) is shorter than an MC simulation running *M_F_* × *N* photons, i.e *T_MC_(N)* + *T_f_* < *T_MC_*(*M_F_* × *N*). Due to space limitations, we are unable to list all combinations of simulations that satisfy the above condition. However, our general observations include 1) the CNN inference runtime is independent of number of simulated photons, 2) DnCNN is typically faster than other CNN denoisers, but also has the poorest performance among them from Table 3, 3) the larger the domain size, the longer it takes for CNN denoisers to run, 4) generally speaking, applying CNN denoisers to simulations with 10^7^ photons or above can result in significant reduction of total runtime.

In our previous work,^16^ we had also concluded that 10^7^ photon is a general threshold for GPU-ANLM to be effective; however, from the runtime data reported here using NVIDIA A100 GPUs, GPU-ANLM appears to also benefit simulations with 10^6^ photons, likely due to the high computing speed of the GPU. Nonetheless, comparing to most tested CNN denoisers, the GPU-ANLM denoiser offers dramatically less equivalent acceleration despite its fast speed.

## 4 Conclusion

In summary, we have developed a framework for applying state-of-the-art DL methods for denoising 3-D images of MC photon simulations in turbid media. A list of supervised CNN denoisers, including DnCNN, UNet, ResMCNet, and DRUNet, were implemented, extended for processing 3-D data, and tested for denoising MC outputs. In addition, we have developed a customized cascaded DnCNN/UNet denoiser combing the global-noise removal capability of DnCNN and local-noise removal capability of UNet. All developed MC denoising networks were trained by using GPU accelerated MCX simulations of random domains to learn the underlying noise from MC outputs at a range of photon numbers. A simple yet effective synthetic training data generation approach was developed to produce complex simulation domains with random inclusions made of of 3-D polyhedral and ASCII characters with random optical properties and simulation parameters. In addition to following current best practices of contemporary CNN and DL development, we have also specifically fine-tuned and customized our MC denoisers to better handle the unique challenges arose in denoising 3-D MC data. For example, to handle the high dynamic range in MC fluene maps using CNNs, a reversible log-mapping scheme was applied to each volume before being fed to the models. In addition, we have also applied inference twice and combined the results to further enhance the dynamic range of the input data. All reported CNN MC denoisers have been implemented in the Python programming language using the PyTorch framework, with both source codes and training data freely available to the community as open-source software.

To evaluate the efficacy of these proposed CNN denoisers, we have constructed 6 standard benchmarks – 3 simple domains and 3 complex ones – from which we have derived and reported both global performance metrics (such as SSIM and PSNR) and local performance metrics (such as Δ*SNR* and *M_F_*). From our results, all tested CNN based denoisers offered significantly improved image quality compared to model-based image denoisers such as GPU-ANLM and BM4D in this particular application. Overall, most CNN denoisers provide a 10 to 20 dB SNR improvement on average, equivalent to running 10 to 100 fold more photons. Among these CNN denoisers, our proposed Cascade network outperformed most of the state-of-the-art spatial domain denoising architectures and yielded the best image quality for low-photon simulations with 10^5^ and 10^6^ photons. Its performance is on-par or only slightly inferior compared to DRUNet in high-photon simulations (10^7^ photon) in simple domain tests. For all benchmarks involving real-world complex domains, the Cascade network yielded the highest global metrics in nearly all tests. In comparison, some of the most effective model-based image denoisers such as the GPU-ANLM filter we proposed previously^16^ only yielded 3-4 dB improvement, despite being relatively fast to compute. It is worth noting that the Cascade network yielded an impressive 80-fold equivalent speedup when processing low-image-feature simulations such as a homogeneous domain.

From our tests, CNN denoisers demonstrate superior scalalability to input data sizes and input image qualities. Although our training data were produced on a 64 × 64 × 64 voxelated space with relatively simple shapes, all tested CNN denoisers show no difficulty in handling images of larger sizes or significantly more complex inclusions. Our Cascade network also reported a 12 dB average SNR improvement when being applied to denoise baseline simulations with 10^9^ photons – the level of photon number that was used as the “ground-truth” for training. This suggest that these CNN denoising architectures may not be strictly limited by the quality of the data that they are trained on.

From our results on runtimes, most CNN denoiser inference (including two passes) time ranges between less than a second to a dozen seconds, regardless of the input data quality. We concluded that in order to yield an overall shorter total runtime, applying CNN denoisers to processing MC images generated from 10^7^ photons or more can generally lead to significantly improved computational efficiency.

One of the limitations of the current work is the relatively long training time. To train each denoising network using our synthetic dataset of 1,500 random domains (each with 5 photon number levels with multiple rotated views) requires on-average a full day (24 hours) if running on a high-end 8-GPU server with large-memory NVIDIA A100 GPUs (40 GB memory allows to use a batch-size of 4 for acceleration). If running on a single GPU node, we anticipate the required training time is around 10 to 12 days on a single A100 GPU, and even longer for low-memory GPUs. Experimenting with the number of layers in each model to reduce the number of intermediate tensors while retaining the performance benefits reported in this work, as well as the development of new and significantly more compact deep-learning based denoisers will be the focus of our future work. Moreover, some of the training parameters were determined empirically and deserve further optimization. For example, we trained the networks over 64 × 64 × 64 domains. It could be significantly faster if we can reduce the training data size while still retain the scalability to arbitrarily sized domains. Additionally, the landscapes of CNN architecture and denoising networks are constantly being updated and improved over the past few years. We can not exhaust all emerging CNN denoisers and would be happy to extend this work with newer and more effective CNN denoising architectures in the future.

To conclude, we strongly believe that investigating high-performance image denoising techniques offers a new direction for researchers seeking for the next major breakthrough in speed acceleration for MC simulations. DL and CNN based image denoising techniques have demonstrated impressive capabilities compared to the more traditional model-based denoising methods, and yielded notable image quality enhancement that is equivalent to running 10 to 50 times more photons, which can be directly translated to 10 to 50 fold speedup, in most of our tested benchmarks. Our Cascade denoising network even reported a nearly 80-fold equivalent speedup when denoising homogeneous domain results – a level of acceleration we were only able to witness when migrating MC from single-threaded computing to massively parallel hardware over a decade ago.^11–13^ With the combination of advanced image processing methods and new simulation techniques, we anticipate MC to play an increasingly important role in today’s biomedical optics data analysis and instrument development. All software and data generated from this work, including our Python implementations of various CNN denoisers and scripts to recreate the training/testing datasets, are freely available to the research community as open-source software and can be downloaded at http://mcx.space.

## Acknowledgments

This research is supported by the National Institutes of Health (NIH) grants R01-GM114365, R01-CA204443 and R01-EB026998. LY acknowledges the insightful inputs and discussions related to UNet from Dr. Hang Zhao at MIT Media Lab. This work was completed in part using the Discovery cluster, supported by Northeastern University’s Research Computing team.

